# Genome-Wide Identification and Characterization of *MLO* Gene Family in Octoploid Strawberry (*Fragaria ×ananassa*)

**DOI:** 10.1101/2020.02.03.932764

**Authors:** Ronald R. Tapia, Christopher R. Barbey, Saket Chandra, Kevin M. Folta, Vance M. Whitaker, Seonghee Lee

## Abstract

Powdery mildew (PM) caused by *Podosphaera aphanis* is a major fungal disease in cultivated strawberry. *Mildew Resistance Locus O* (*MLO*) is a gene family described for having conserved seven-transmembrane domains. Induced loss-of-function in specific *MLO* genes can confer durable and broad resistance against PM pathogens. However, the underlying biological role of *MLO* genes in strawberry is still unknown. In the present study, the genomic structure of *MLO* genes were characterized in both diploid (*Fragaria vesca*) and octoploid strawberry (*Fragaria ×ananassa)*, and the potential sources of *MLO*-mediated susceptibility were identified. Twenty *MLO*-like sequences were identified in *F. vesca*, with sixty-eight in *F. ×ananassa*. Phylogenetic analysis divides strawberry *MLO* genes into eight different clades, in which three *FveMLO* and ten *FaMLO* genes were grouped together with the functionally known *MLO* susceptibility. Out of ten *FaMLO* genes, *FaMLO17-2* and *FaMLO17-3* showed the highest similarity to the known susceptibility MLO proteins. Gene expression analysis of *FaMLO* genes was conducted using a multi-parental segregating population. Three expression quantitative trait loci (eQTL) were substantially associated with *MLO* transcript levels in mature fruits, suggesting discrete genetic control of susceptibility. These results are a critical first step in understanding allele function of *MLO* genes, and are necessary for further genetic studies of PM resistance in cultivated strawberry.

## Introduction

Powdery Mildew (PM) is a major fungal plant disease affecting many economically important cereal and horticultural crops, including strawberry. The causal agent of PM in strawberry is the obligate parasite *Podosphaera aphanis* (Wallr.) (former *Sphaerotheca macularis* f. sp. *fragariae*)^1^. PM primarily affects the leaf and depending on severity can also affect other organs above ground^2^. Initially, a white powdery mycelium develops on the underside of the leaves, followed by leaf upward curling while severe leaf infection can cause burning at leaf margins^3^. Infected flower and fruits, on the other hand, can cause fruit deformation and delayed ripening^4^. PM is widespread in many strawberry growing regions worldwide, such that either open-field or high-tunnel growing system can experience severe yield losses when infected field are left untreated^5^. Most farmers rely heavily on multiple antimicrobial chemistry applications to manage PM in the field. Hence, developing new cultivars with improved resistance against powdery mildew is necessary to significantly reduce reliance to anti-fungal applications and promote sustainable agriculture.

The *Mildew Resistance Locus O* (*MLO*) gene family is present in several crop species and was described for having conserved seven-transmembrane (TM) and C-terminal calmodulin-binding (CaMB) domains^6,7^. The *MLO* gene family has been the focus of attention in many crop species because transgenic downregulation or elimination of specific endogenous *MLO* genes has led to PM resistance^8^. However, alteration of *MLO* gene sequences can trigger negative phenotypic effects including premature leaf chlorosis, altered root growth and pollen tube germination^9^.

The first *MLO*-based resistance trait was first characterized in barley (*HvMLO*), where a loss-of-function mutation in an *MLO* gene conferred broad resistance against PM pathogens^10^. This discovery led to subsequent comparative genomic studies of *MLO* gene families in several plant species to find suitable candidate genes. In the model plant *Arabidopsis thaliana*, three *MLO* genes (*AtMLO2*, *AtMLO6* and *AtMLO12*) were functionally characterized to confer PM susceptibility^11,12^. Expression analysis of these homologs upon pathogen challenge suggested functional redundancy. The *Atmlo2* single mutant has only partial resistance while triple mutants (*Atmlo2, Atmlo6* and *Atmlo12*) have full resistance against PM pathogens^13^. Identification of the functional *MLO* genes greatly relies on the availability of genetic information, therefore the development of reference genomes for several crop species provides an opportunity to identify potential *MLO* orthologs which can be used for functional studies.

*MLO* genes have been genetically characterized across many crops including apple (*MdMLO1*), pepper (*CaMLO2*), rose (*RhMLO1)*, grapevine (*VvMLO13*), melon (*CmMLO2*), pea (*PsMLO1*), tobacco (*NtMLO2*), tomato (*SlMLO1*), rice (*OsMLO1*), corn (*ZmMLO1*) and wheat (*TaMLO1*)^8,14,15^. Recently, targeted-genome mutation of the *SlMLO1* gene resulted in the development of PM-resistant *Slmlo1* tomato variety^16^. The durable and broad-spectrum effectiveness of *MLO*-based resistance could provide a useful target for strawberry improvement to PM disease.

The modern cultivated strawberry (*Fragaria ×ananassa*) is an allo-octoploid (2n=8x=56) resulting from hybridization between a Chilean strawberry (*F. chiloensis*) and a North American native strawberry (*F. virginiana*)^*17*^. Further domestication of *F. ×ananassa* produced large and tasty berries that have become one of the world’s most widely grown fruit crops. In 2010, the genome of the diploid progenitor species *F. vesca* was sequenced^18^ and has been used as an excellent genetic resource towards molecular marker development^19^ and for gene-trait association studies in *Fragaria* species^20–22^. However, the *F. ×ananassa* genome is far more complicated than its diploid progenitor, as diploid species contain two copies for each gene whereas octoploid species may possess up to eight. Recently, a new *F. ×ananassa* reference genome was developed, and the complex evolutionary origin of octoploid strawberry has been deciphered^23^. This new reference genome sequence will serve as powerful genetic resource to unravel complexity of octoploid strawberry genome for gene-traits association studies including causal *MLO* genes in strawberry breeding programs.

The goals of this study were to characterize *MLO* gene family in diploid (*F. vesca*) and octoploid (*F. ×ananassa*) strawberries, compare their gene structures, compare transcript levels throughout the plant and between individuals, and identify potential *MLO* genes associated with PM resistance and fruit ripening in octoploid strawberry. The data reported here are a critical first step in understanding allele function of strawberry *MLO* genes which will be useful for future genomics and functional studies.

## Results

### Genome-wide identification of *MLO* gene family in diploid and octoploid strawberry

Using the Arabidopsis MLO protein sequence as a BLAST query, 20 putative *MLO* genes were identified using the latest diploid genome annotation *F. vesca* v4.0.a1^24^. These 20 *MLO* genes were renamed *FveMLO1* through *FveMLO20* based on their ordered chromosomal positions (Fig. 1A, Table S1a). Predicted proteins ranged between 200 to 903 aa with an average of 524 aa. Three truncated MLO proteins were identified as putative pseudogenes (Table S2). The number of predicted *MLO* genes in diploid strawberry agrees with the previous characterization of strawberry *MLO* genes using the first *F. vesca* draft genome^25^.

The *FveMLO* genes identified in *F. vesca* were then used to identify gene orthologs in the octoploid strawberry annotated genome *F. ×ananassa* v1.0.a1^23^. Analysis revealed 68 predicted *MLO* genes across 28 chromosomes (Fig. 1B, Table S1b). Of 20 *FveMLO* genes, 17 were matched with high amino acid sequence identity to a putative ortholog, however, conserved sequences could not be found for *FveMLO6*, *FveMLO7* or *FveMLO8*. Based on their putative orthology to *F. vesca*, the octoploid *MLO* genes were named *FaMLO1* through *FaMLO20,* respectively. To distinguish homoeologous *MLO* genes, *FaMLO1-1 to FaMLO1-4* were used for each *FveMLO1* homolog. Predicted MLO proteins have amino acid sequence lengths ranging between 144 and 2,365 with an average of 560 amino acids (Table S2).

**Figure 1.**
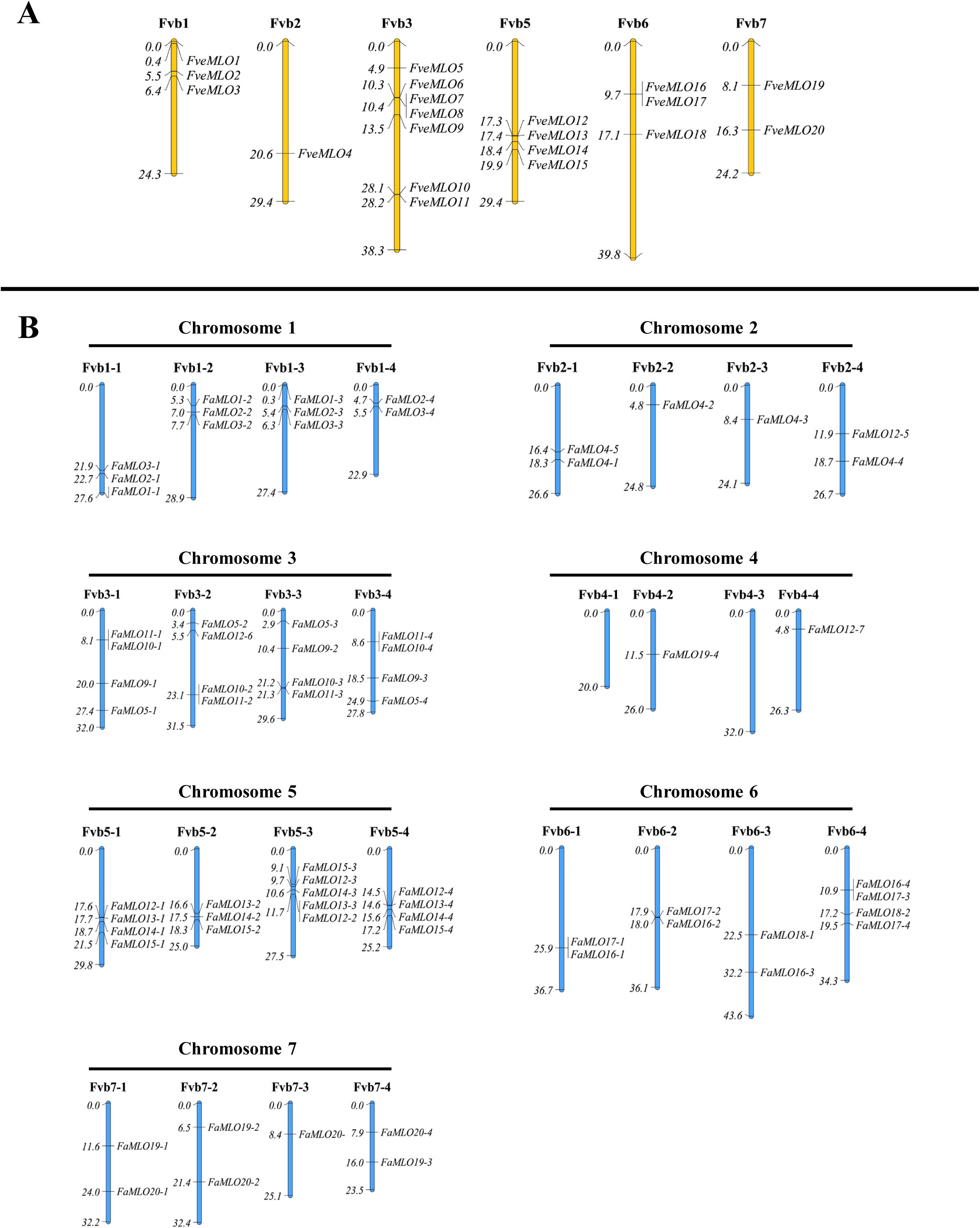
Chromosomal localization and distribution of *FveMLO* (**A**) and *FaMLO* (**B**) genes in *F. vesca* and *F. ×ananassa* genomes, respectively. The seven chromosomes of *F. vesca* (yellow) were named as Fvb1 to Fvb7 while *F. ×ananassa* (blue) were named as Fvb1-1 to Fvb1-7, Fvb2-1 to Fvb2-7, Fvb3-1 to Fvb3-7 and Fvb4-1 to Fvb4-7, respectively to indicate four subgenomes within each chromosome. Putative strawberry *MLO* genes chromosome locations were visualized using MapChart 2.3 software^41^. Relative chromosome size was indicated by unit, Mbp.

### Gene structure and synteny analysis of octoploid strawberry *MLO* genes

Twenty *FveMLO* genes were distributed randomly across six chromosomes of the diploid *F. vesca* genome: one in Fvb2, two in Fvb7, four in Fvb5, seven in Fvb3 and three in Fvb1 and Fvb6 (Fig. 1, Table S1a). The coding DNA sequence composition ranged from 1 to 16 exons (Fig. S1-A). In the octoploid genome, 68 *FaMLO* genes were distributed across every chromosome except Fvb4-1 and Fvb4-3 (Fig. 1B, Table S1b). The intron-exon structures of *FaMLO* genes have more variation as compared with its diploid progenitor *F. vesca,* with coding DNA sequence (CDS) composition ranging from 1 to 23 exons (Fig. S1-B). Despite this structure complexity, many *FaMLO* genes demonstrated high homology of DNA sequence with the diploid progenitor, *F. vesca* (Fig. S2). Two additional *FaMLO* sequences in chromosome 4-2 (*FaMLO19-4*) and 4-4 (*FaMLO12-7*) were identified in *F. ×ananassa,* which are not present in *F. vesca* genome (Fig. 1A-B).

The four subgenomes in the modern cultivated strawberry are a result of allo-polyploidization with four specific diploid progenitor genomes, which have subsequently undergone substantial subgenome conversion^23^. To elaborate on this recent discovery, we display synteny networks for putative *MLO* genes between diploid (*F. vesca*) and octoploid (*F.×ananassa*) strawberry (Fig. 2). Most putative *FaMLO* orthologs co-localized in the same chromosome of *F. vesca* genome. Additional *FaMLO* orthologs of *FveMLO12* and *FveMLO19* were identified in subgenomes Fvb4-4 and Fvb4-2, respectively. Meanwhile, no orthologs were found for *FveMLO5, FveMLO6* and *FveMLO7* in the *F. ×ananassa* genome. Variants in copy number of homoeologous genes for each *FaMLO* were identified with two homeologs each for *FaMLO18* and *FaMLO19*, three for *FaMLO1* and *FaMLO9,* four for *FaMLO2, FaMLO3, FaMLO5, FaMLO20* and *FaMLO10* through *FaMLO17*, and five for *FaMLO4* (Fig. 2. The variation in the distribution of *MLO* genes showed a wide diversity in the genome composition of octoploid strawberry. Furthermore, there were at least two *FaMLO* homeologs with 80% or more sequence similarity with *FveMLO* orthologs (Fig. S2).

**Figure 2.**
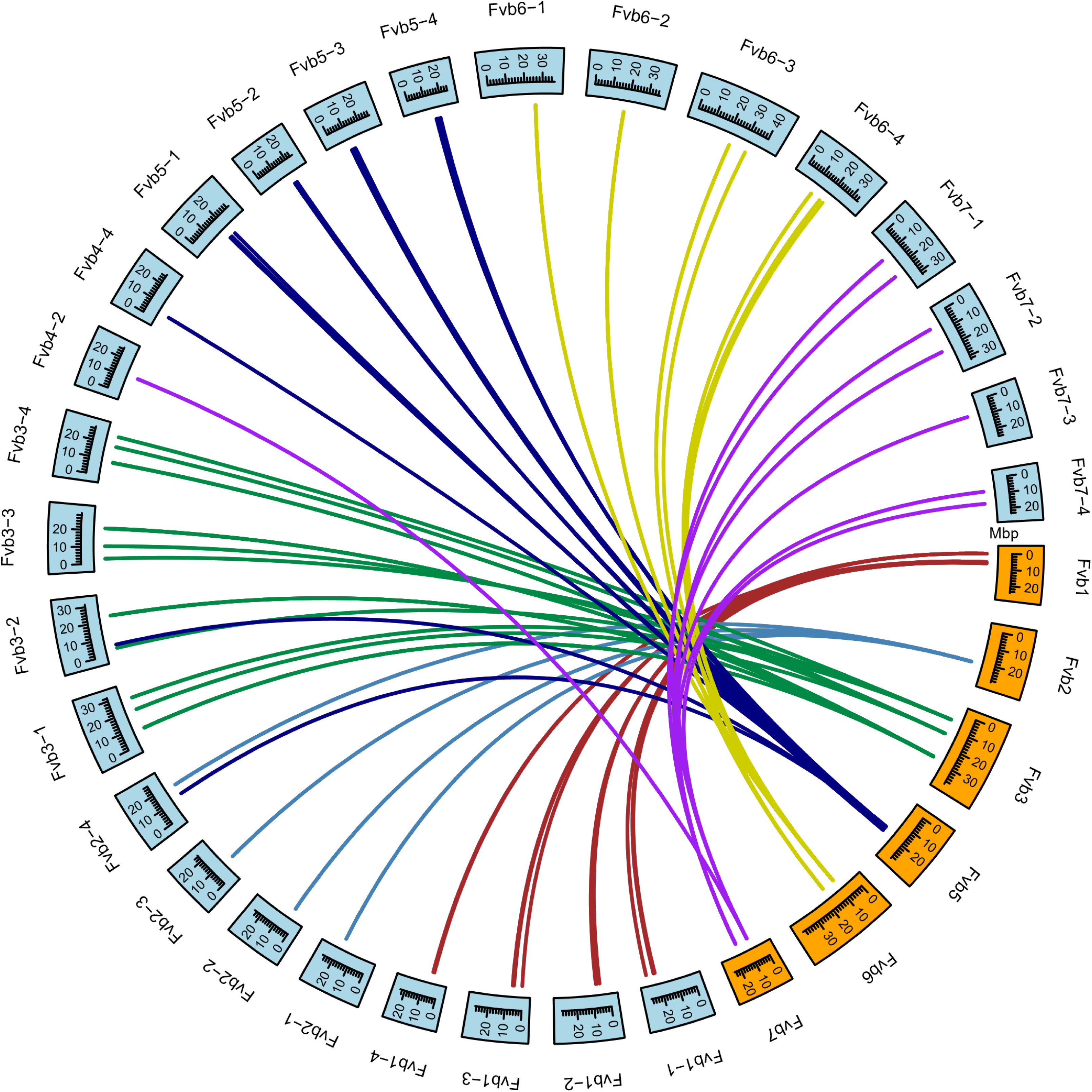
Synteny analysis of *MLO* genes between *F. vesca* and *F. ×ananassa*. Syntenic regions present in each chromosome of *F. vesca* was filled with red, light blue, green, dark blue, yellow and pink sequentially. A total of 68 connecting lines between two genomes denote syntenic chromosomal regions. The *F. vesca* and *F. ×ananassa* chromosomes are highlighted in orange and light blue. Chromosomes Fvb4 from *F. vesca,* and Fvb4-1 and Fvb4-3 from *F. ×ananassa* were not included since no putative *MLO* genes was identified in those regions. Relative chromosome size was indicated by unit, Mbp. Circular visualization of syntenic regions between *F. vesca* and *F. ×ananassa* were constructed using an R-package “Circlize”.

### Domain organization and structure characterization of octoploid strawberry *MLO* genes

The *MLO* gene was first identified in barley and characterized as membrane protein with seven transmembrane (TM) domains and a uniquely-identified “*MLO*-functional domain”^6^. To examine the conserved protein domains of the strawberry *MLO* genes, the deduced amino acid sequence of predicted MLO proteins found in diploid and octoploid strawberries were subjected to theoretical domain prediction using online software InterProScan (https://www.ebi.ac.uk) and NCBI’s conserved domain database (https://www.ncbi.nlm.nih.gov/Structure/cdd). Most MLO proteins from either diploid or octoploid strawberry contain the conserved domain of MLO and TM, which covers a large portion of the protein (Fig. 3). To predict TM domains and subcellular localization of strawberry MLO proteins, CCTOP^26^ and WoLF PSORT^27^ software were used to find differences in the number of TM domains and subcellular localization among FveMLO and FaMLO proteins (Fig. 3, Table S2). All FveMLO proteins were predicted to localize within the plasma membrane except for FveMLO7, which was predicted to localize in the extracellular matrix. Out of 68 *FaMLO* genes, 61 were predicted to localize within plasma membrane while nine were predicted to localize in other organelles: four in the chloroplast, two in ER, two in the nucleus and one in Golgi bodies (Table S2). Thirteen out of 20 FveMLO proteins have seven TM domains, while seven have between three to six TM domains. The *FaMLO* gene family has a high degree of variation in TM domain composition (Table S2). Only 35 FaMLO proteins have seven TM domains while the remaining FaMLO proteins have TM domains ranging between zero and eight (Fig. 3).

**Figure 3.**
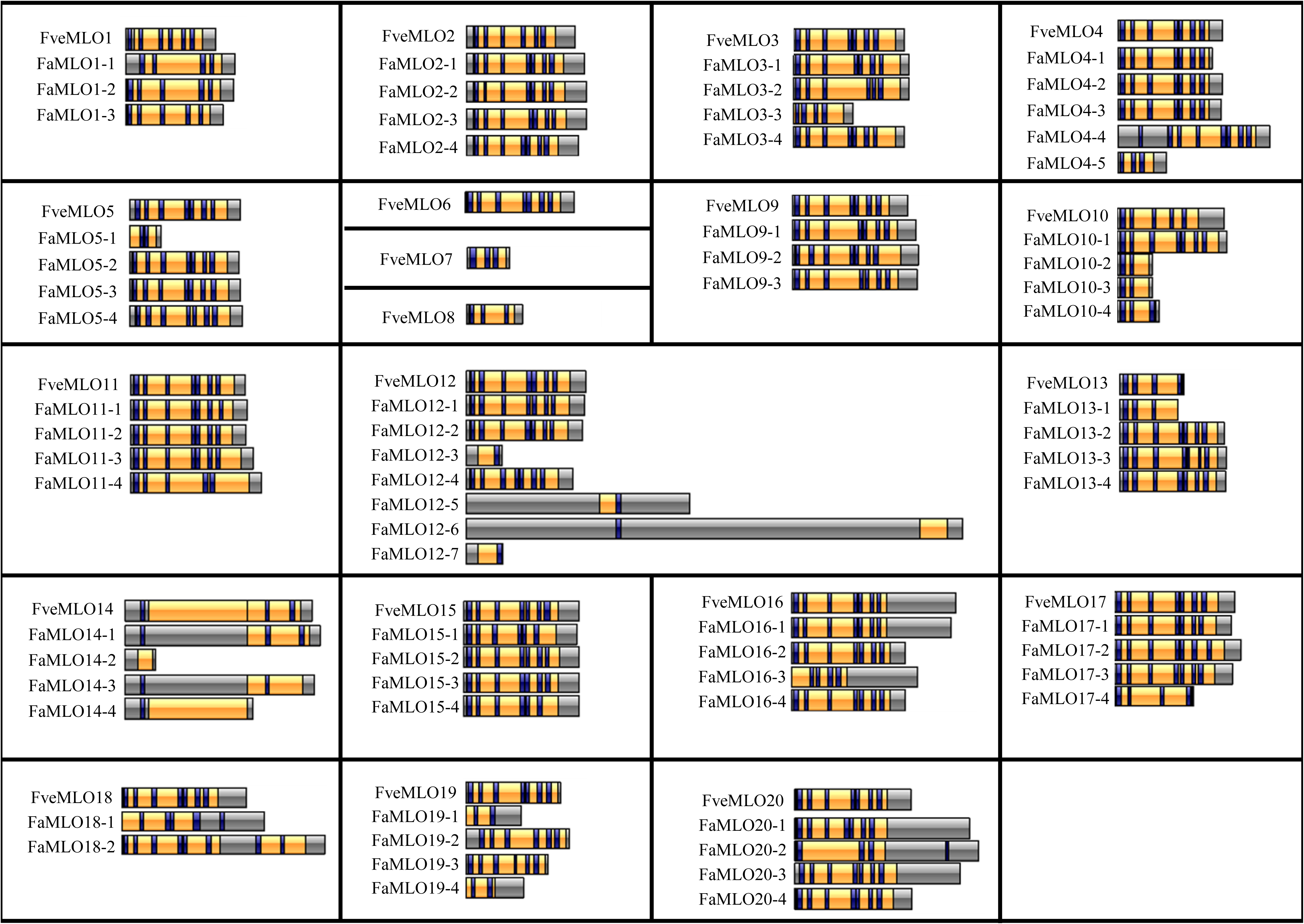
Domain organization of deduced *MLO* protein sequences of *F. vesca* and *F.×ananassa*. Visualization of protein domains was constructed using IBS 1.0.3 (Illustrator for Biological Sequences) program^47^. The positions of transmembrane (blue) and *MLO* (yellow) domains were predicted using online CCTOP prediction server (http://cctop.enzim.ttk.mta.hu/) and CDD: NCBI’s conserved domain database (https://www.ncbi.nlm.nih.gov/Structure/cdd).

The domain organization of some FaMLO proteins showed high levels of conservation with the diploid ancestor species, *F. vesca*. For example, there were at least two homoeologs of *FaMLO2, FaMLO4, FaMLO5, FaMLO9, FaMLO11, FaMLO12, FaMLO15, FaMLO16, FaMLO17* and *FaMLO20* that showed *F. vesca*-like domain structures (Fig. 3). However, one to three homoeologs of some FaMLO proteins including *FaMLO3, FaMLO4, FaMLO5, FaMLO10, FaMLO 12, FaMLO13, FaMLO414, FaMLO16, FaMLO17, FaMLO18, FaMLO19* and *FaMLO20* showed more diverse protein structures which caused by either protein sequence truncation or extension (Fig. 3).

Variation in protein characteristics of *FaMLO* gene family, especially between homoeologous proteins, could imply that some MLO proteins are unique and might not share the same function. MLO proteins that were localized in the plasma membrane and contain conserved transmembrane protein domains were selected for further phylogenetic analysis of functional divergence.

### Phylogenetic relationship analysis of *MLO* genes in strawberry

To study the evolutionary relationships between strawberry *MLO* genes and *MLO* genes from other plant species, we aligned the deduced amino acid sequence of 20 *MLO* genes from *F. vesca* (*FveMLO*s) and 68 from *F. ×ananassa* (*FaMLOs)* with previously characterized *MLO* from Arabidopsis (*AtMLO*s), rice (*OsMLO1*), corn (*ZmMLO1*), barley (*HvMLO*), tomato (*SlMLO1*), pepper (*CaMLO2*) and other rosaceous crops such as apple (*MdMLO*s) and peach (*PpMLO*s). Phylogenetic analysis using MUSCLE and FastTree divided *MLO* genes into eight different clades (Fig. 4). Functionally characterized *MLO* genes that are associated with PM resistance from selected monocots and dicots were clustered in clade IV and V, respectively. Among strawberry *MLO* genes, *FveMLO12* and *FaMLO12* together with *HvMLO, OsMLO1* and *ZmMLO1* were grouped in clade IV while *FveMLO10*, *FveMLO17*, *FveMLO20, FaMLO10, FaMLO17* and *FaMLO20* together with *SlMLO1, CaMLO1, AtMLO2, AtMLO6* and *AtMLO12* were grouped in clade V. Remaining putative *MLO* sequences were distributed to six other groups. The results showed that three *FveMLO* and ten *FaMLO* proteins have a close evolutionary relationship with the MLO proteins known to confer PM susceptibility.

**Figure 4.**
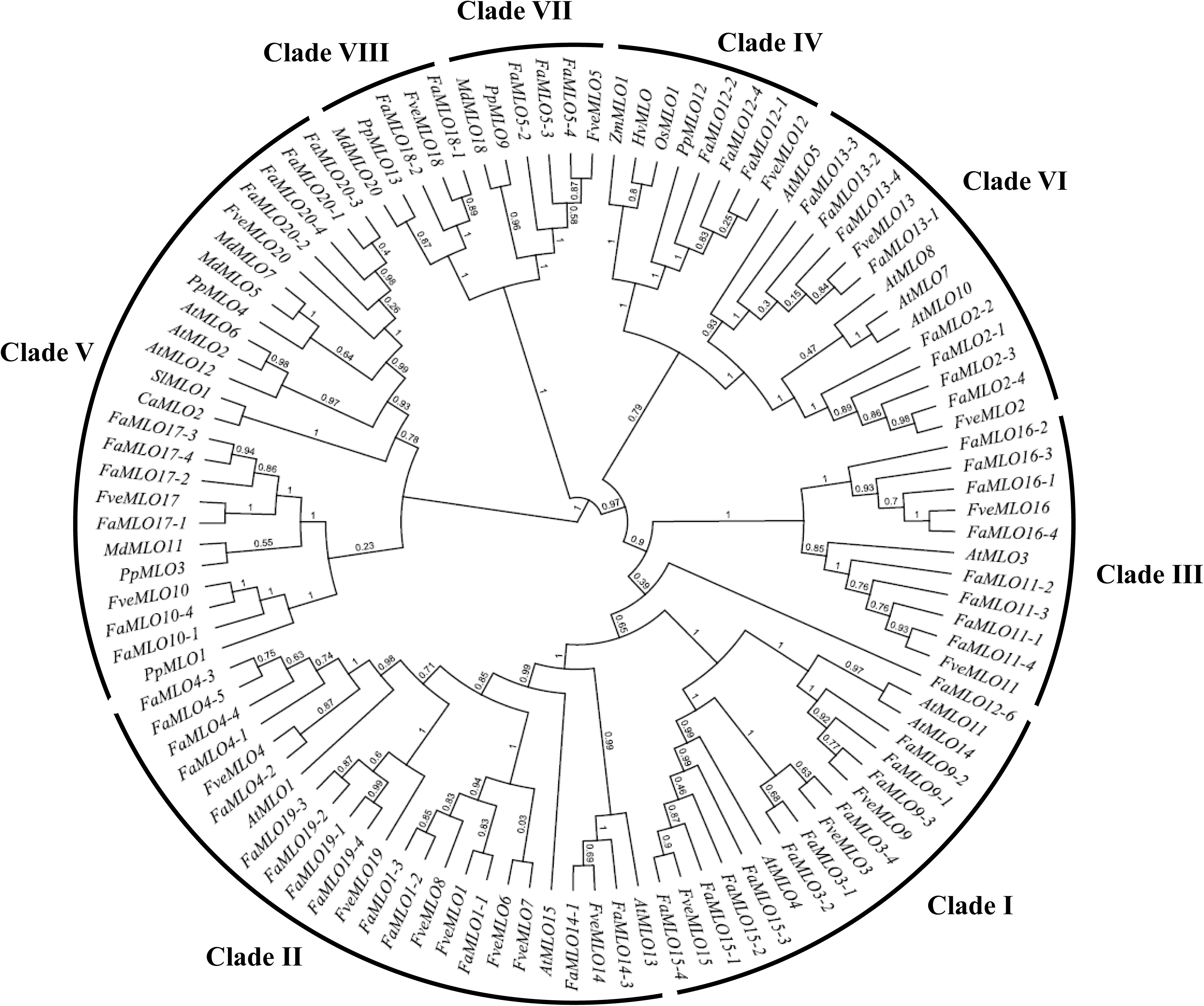
Phylogenetic tree of *MLO* protein sequences of strawberry., *F. vesca* and *F.×ananassa,* and other plant species such as Arabidopsis, tomato, pepper, rice, barley, maize, apple and peach. Strawberry *MLO* genes together with other *MLO*s were grouped into eight different clades. Unrooted phylogenetic tree of 15 *AtMLOs,* 20 *FveMLO*s, 59 *FaMLOs,* five *PpMLO*s, six *MdMLO*s, *CaMLO2, SlMLO1, HvMLO, OsMLO1* and *ZmMLO1* was constructed by using FastTree via Geneious software.

To further investigate the candidate susceptibility-conferring *FaMLO* genes, namely *FaMLO10-1*, *FaMLO10-4*, *FaMLO17-1*, *FaMLO17-2*, *FaMLO17-3*, *FaMLO17-4*, *FaMLO20-1*, *FaMLO20-2*, *FaMLO20-3* and *FaMLO20-4,* their deduced amino acid sequences were aligned with their orthologs from *F. vesca* and *AtMLO2, AtMLO6* and *AtMLO12* from Arabidopsis (Fig. S3). We identified conserved domains including seven TM, CaMB and two C-terminal protein (I and II) domains of *MLO* genes^7,28^. *FaMLO17-2*, *FaMLO17-3*, *FaMLO20-1*, *FaMLO20-2*, *FaMLO20-3* and *FaMLO20-4* proteins possessed seven TM domains and conserved CaMB and C-terminal I and II domains; however, *FaMLO10-1, FaMLO10-4, FaMLO17-2* and *FaMLO17-4* showed truncated protein sequences at C-terminal end, resulting in the loss of these conserved domains. Among candidate *MLO* proteins, *FaMLO17-2* and *FaMLO17-3* protein sequences are most identical to known susceptibility-conferring Arabidopsis *MLO* proteins, and therefore could have potential association with PM resistance in octoploid strawberry.

### Expression profiling of *FaMLO* genes in cultivated strawberry

Expression patterns of previously characterized *MLO* genes suggested diverse biological functions for the *MLO* gene family^29^. To investigate functional strawberry *MLO* genes in fruit, we performed RNA sequencing analysis to evaluate transcript profiles in selected strawberry varieties, ‘Mara de Bois’, ‘Florida Elyana’, ‘Winter Dawn’, ‘Florida Radiance’ and ‘Strawberry Festival’. Variation in transcript abundance was observed among *MLO* genes and 27 *FaMLO* genes were constitutively expressed (>1 Transcript per Million, TPM) across samples (Fig. 5A and Table S3). We found that *FaMLO14* homeologs had the highest fruit transcript abundance among *FaMLO* genes. Meanwhile, *FaMLO1, FaMLO13 and FaMLO15-18,* including their homeologs, were not detected in all varieties (Table S3). *MLO* genes clustered in clade V also showed variation in gene expression among them and between their homoeologous genes. Transcripts from at least two homeologs of *FaMLO10, FaMLO17* and *FaMLO20* were detected suggesting that genes are functional. To characterize *MLO* transcript accumulation patters in octoploid strawberry, raw data from the strawberry gene expression atlas study^30^ were re-assembled using the recently published octoploid strawberry genome^23^. The steady-state accumulation of several *MLO* transcripts are specific to different tissues, with *MLO* genes such as *FaMLO1* and *FaMLO17* showing tissue-specific gene expression. The gene *FaMLO17* and its two homeologs are preferentially expressed in root tissues (Fig. 5B and Table S4).

**Figure 5.**
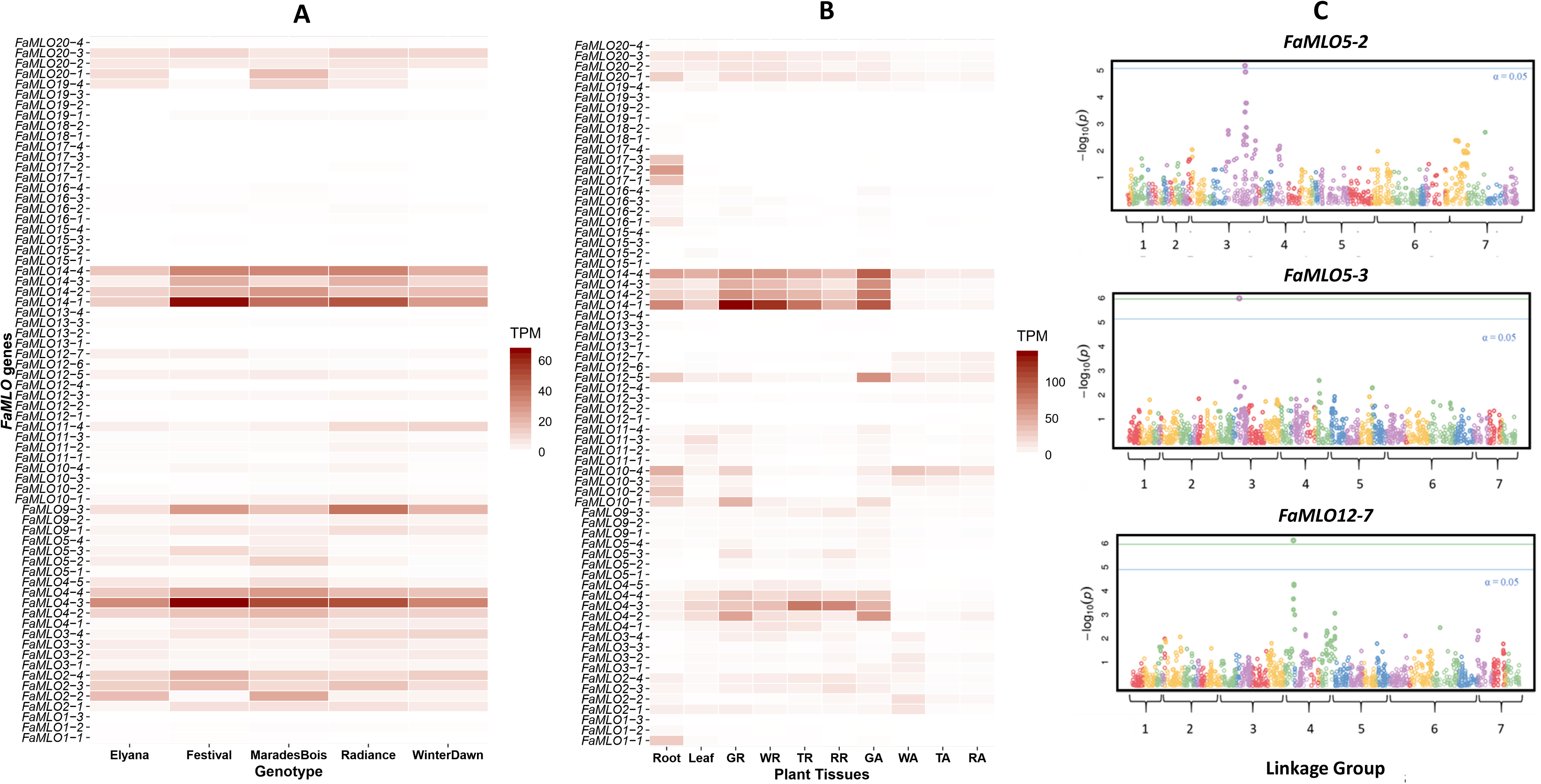
Transcript and GWAS analysis of *FaMLO* genes. A) Expression profile of putative strawberry MLO genes in mature fruit of selected strawberry varieties, ‘Mara de Bois’, ‘Elyana’, ‘Winter Dawn’, ‘Radiance’ and ‘Festival’. B) Expression profile in different plant tissues e.g., leaf, roots, green (GR), turning (TR), white (WR) and red (RR) receptables, and green (GA), turning (TA), white (WA) and red (RA) achenes extracted from RNA-seq data published by Sánchez-Sevilla, et al (2017). The mean gene expression level was normalized using Transcripts Per Million (TPM). Gene expression profile was visualized with heatmap using R-package “ggplot2”. C) Manhattan plots of GWAS showing significant SNPs associated with *FaMLO* transcript in fruits. X-axis corresponds to the distribution of SNPs across seven chromosomes of *F. vesca* while y-axis shows FDR-adjusted probability values.

### Identification of eQTL associated with *FaMLO* transcripts in strawberry mature fruit

RNA-seq analysis identified *FaMLO* transcript level variation across fruit developmental stages (Fig. 5B), supporting a potential role of *MLO* gene family in the fruit. To characterize *FaMLO* fruit expression variation associated with genetic differences between individuals (eQTL), we performed eQTL analysis using 61 octoploid IStraw35 genotypes and mature fruit transcriptomes and identified at least three eQTL. These include maker-Fvb4-4-augustus-gene-48.50 (*FaMLO12-7*), snap_masked-Fvb3-2-processed-gene-33.29 (*FaMLO5-2*) and snap_masked-Fvb3-3-processed-gene-28.38 (*FaMLO5-3*), the latter two of which are homoeologous genes (Fig. 5C). Single marker analysis of *MLO* eQTL also shows a general association between homozygous marker genotypes and lower *MLO* transcript levels (Fig. S4). Because the ‘Camarosa’ genomic locus of each transcript is known, it could be shown that some of the identified transcript eQTL coincide with the location of their corresponding *MLO* gene locus, and hence were considered as cis-eQTL (Table 1). The most significant IStraw35 SNP marker name and position for each *MLO* gene is provided with the eQTL phase, minor allele frequency, p-value (FDR-adjusted) and heritability estimates (Table 1).

**Table 1.** Summary genome-wide association analysis of loci significantly associated with *MLO* transcript in strawberry fruit. The most significant IStraw35 SNP marker name and position for each *MLO* gene is provided with the eQTL phase, minor allele frequency, p-value (FDR-adjusted), and heritability estimates.

## Discussion

The *MLO* gene family is an important target in many agricultural crops for the improvement of resistance against PM pathogens. The previous characterization identified that some *MLO* genes are involved in PM susceptibility, and that loss-of-function of those genes can confer durable and broad-spectrum resistance. In this study, it was found that variants of *F. vesca MLO* genes and *F. ×ananassa MLO* genes were distributed across the genomes. However, we observed copy number variation of *FaMLO* homeologs showing differences in composition among subgenomes, supporting the distinct origin of each subgenome during the evolution of octoploid strawberry^29^. The unique *FaMLO* sequences in chromosome four (Fig. 1B and Fig. 2) that are not present in *F. vesca* might suggest an acquisition of chromosome segments from other diploid progenitors. Also, we did not find any significant homologs for *FveMLO6, FveMLO7* and *FveMLO8* using the *F. ×ananassa* ‘Camarosa’ reference genome (Fig. 1B). The loss of genes could be evolutionary significant in the development of polyploid species^31^ or other possible functional divergence during domestication. Additional octoploid reference genomes would explain such phenomena, as the chromosomal localization of *FaMLO* genes and possible misorientation in genome sequence for some subgenomes can be identified.

The average MLO protein length (~500 amino acids) was similar in both *F. vesca* and *F.×ananassa*, although few *FaMLO* proteins had more structural variations (SV) as compared with other homologous genes (Fig. 3). SV is defined as a change in genome sequence such as deletion, insertion and inversion caused by mutations which in turn can affect biological gene function^32^. Furthermore, SV between homeologs of *FaMLO10* and *FaMLO12* causes changes in their gene expression profile (Fig. 5A-B) and subcellular localization (Table S2). SV between homeologs might result in different gene regulation of gene expression and consequently lead to novel functions^33^. Diverse gene expression patterns among *FaMLO* genes across different tissues (Fig. 5B) suggested that MLO proteins might be specialized for different biological responses. This may be analogous to the unique expression patterns observed for Arabidopsis *MLO* genes, where transcription of each *MLO* gene is distinct and regulated differently by various biotic and abiotic stimuli^34^

In the present study, we found that number of the TM domains of *FveMLOs* and *FaMLOs* varied between zero to eight. This variation was also observed in previous genome-wide *MLO* studies in other Rosaceous crops such as apple and peach^35^. Phylogenetic analysis grouped strawberry *MLO* genes into eight different clades, which is consistent with previously classified *MLO* gene families in diploid strawberry^25^ and other Rosaceous crops^35^. Functional *MLO* genes associated with PM resistance are grouped in clade IV and V for monocots and dicots, respectively^13^. Interestingly, we found *FveMLO12*, *FaMLO12-1, FaMLO12-2* and *FaMLO12-4* grouped together in clade IV with previously characterized *HvMLO, OsMLO*, *ZmMLO1* and *PpMLO12* ^35^. In the present study, we found at least three clade V *FveMLO* and *FaMLO* genes, which are potential candidate genes involved in PM resistance. *MLO*-based resistance to the PM pathogen *P. aphanis* in a strawberry was successfully tested through RNAi-induced gene silencing of *FaMLO* genes using antisense *MLO* from peach (*PpMLO1*)^36^. In Arabidopsis, mutation of *AtMLO2* showed only partial resistance while complete resistance was obtained after knocking-out both *AtMLO6* and *AtMLO12*^37^. Redundancy of gene function among homologs could arise from gene duplication for adaptation to varying selection pressures and changing environments^38,39^. High gene copy-number is commonly observed in polyploid crop species due to the presence of homoeologous genes, posing a considerable challenge for determining the biological contribution of specific genes. The genes *FaMLO17-2* and *FaMLO17-3* showed the highest sequence identity with known Arabidopsis *MLO* genes, suggesting that they might contribute to strawberry MLO susceptibility. Overall, we identified a total of 10 *FaMLO* genes with high homology to functional disease-susceptible *MLO*, representing candidate genes for breeding new cultivars with improved PM resistance.

Despite considerable research into *MLO* as a susceptibility gene, the endogenous function of the MLO family remains largely unknown. It appears most likely that specific *MLO* genes have diverse and specialized functions^29^. The diverse pattern of expression for *FaMLO* genes in different fruit developmental stages (Fig. 5B), and the presence of certain signal transduction domains, supports one possible role for *MLO* as a ripening regulator. The conserved CaMB domain in *FaMLO* gene family could be a binding site for CaM-dependent modulation during fruit ripening. Calmodulin (CaM) is a well-characterized protein known as Ca^2+^ sensor that triggers response to different developmental and biotic stimuli^7,40^. A previously-studied CaMB domain-containing gene family in tomato showed fruit-ripening regulation activity, via developmental network interaction with Ca^2+^ and calmodulin^40^. Furthermore, generated ripening mutant altered expression of those genes indicating its important role in tomato fruit development. It is possible that the conserved CaMB domain in strawberry *MLO* gene family, together with CaM and other unknown regulators, could play a similar role in the fruit-ripening process. Findings from this study presented the functional diversity of *MLO* gene family, although further studies are needed to validate the integral role of strawberry *MLO* genes in response to biotic and developmental stimuli.

## Conclusions

Here, we identified a total of 20 *MLO* homologues in *F. vesca* and 68 in *F.×ananassa*. Three *FveMLO* genes and ten *FaMLO* genes were clustered with previously characterized *MLO* genes known to be in PM resistance/susceptibility in other plant species. The deduced amino acid sequences of putative strawberry *MLO* genes showed conserved protein characteristics, including transmembrane and calmodulin-binding domains that have been previously described. The considerable amino acid-level variation between MLO homoeologous copies was observed, suggesting possible non-redundant functions of MLOs in different subgenomes. *FaMLO17-2* and *FaMLO17-3* are the most identical to functionally characterized *MLO* genes associated with PM susceptibility in other plants. These genes could be functionally characterized via CRISPR gene editing in future studies. eQTL analysis also identified MLO genes whose differential transcription is governed by genetic segregation, which could be an indication of functional diversity. These findings support and expand upon the potential roles for *MLO*-regulated fruit development. Taken together, these data are a critical first step in understanding the allele function of straberry *MLO* gene family, and should be useful for future functional studies to better understand their role on powdery mildew resistance in strawberry.

## Materials and Methods

### Identification of the strawberry *MLO* genes in diploid and octoploid strawberry

To identify *MLO* gene orthologs in diploid strawberry, *F. vesca*, Arabidopsis *MLO* genes were searched to the latest *F. vesca* genome (v4.0.a1)^24^. Consequently, each *FveMLO* gene identified was used to search for predicted *MLO* genes in octoploid strawberry using the octoploid strawberry reference genome (*F. ×ananassa* v1.0.a1)^23^. The deduced diploid and octoploid strawberry protein sequences were validated by reciprocal BLAST searches obtained from NCBI data sets of Arabidopsis reference genome.

### Gene structure and synteny analysis of strawberry *MLO* genes

The chromosomal localization of each predicted *MLO* genes in *F. vesca* and *F.×ananassa* was identified using available information in GDR database and visualized using MapChart 2.3 software^41^. Gene structure featuring introns, exons and UTR of predicted *MLO* genes were constructed using Gene Structure Display Server 2.0 (GSDS) (http://gsds.cbi.pku.edu.cn/)^42^. Pairwise comparisons of coding DNA sequence (CDS) between predicted *FveMLO* and *FaMLO* genes were obtained using ClustalW2 multiple sequence alignment via Geneious software^43^ followed by heat map visualization to determine closely related *MLO* genes using R-package “Lattice”^44^. Synteny analysis of *MLO* genes between *F. vesca* and *F. ×ananassa* were summarized using R-package “Circlize”^45^.

### Phylogenetic analysis of strawberry and other plant *MLO* genes

The protein sequences of *MLO* genes from *F. vesca* and *F. ×ananassa* were aligned to available sequences and phylogenetic relationship was constructed using FastTree consensus tree protein alignment via Geneious Tree Builder software^43^. Phylogenetic tree was constructed by adding 15 *MLO* genes from Arabidopsis, six *MLO* from other Rosaceous crops, *Malus domesticus* (*MdMLO*s) and *Prunus persica* (*PpMLOs*), barley (*HvMLO*), corn (*ZmMLO1*), rice (*OsMLO1*), tomato (SlMLO1) and pepper (*CaMLO2*).

### Protein characterization and domain prediction

The deduced amino acid sequences of putative *FveMLO* and *FaMLO* genes were analyzed by different prediction software to identify functional domains and determine protein topologies and sub-cellular localizations. Functional MLO domain of protein sequences was predicted using CDD: NCBI’s conserved domain database (https://www.ncbi.nlm.nih.gov/Structure/cdd)^46^. Protein topology and number of transmembrane domains were predicted using online software CCTOP Prediction Server^26^ while protein sub-cellular localization was analyzed using WoLF PSORT program^27^. Default setting was used to run for all prediction software. Visualization of proteins domains were constructed using IBS 1.0.3 (Illustrator for Biological Sequences) program^47^. To analyze conserved amino acids of *MLO* genes associated with PM resistance, protein sequences of *FveMLO10*, *FveMLO17* and *FveMLO20* from *F. vesca* and *FaMLO10*, *FaMLO17* and *FaMLO20* from *F. ×ananassa* were aligned against functionally characterized *AtMLO2*, *AtMLO6* and *AtMLO12* from Arabidopsis using MultAlin software (http://multalin.toulouse.inra.fr/multalin/)^48^.

### Expression profile of putative strawberry *MLO* genes

To examine transcript accumulation patterns of putative strawberry *MLO* genes, we performed RNA-seq analysis in selected strawberry cultivars including ‘Florida Elyana’, ‘Mara de Bois’, ‘Florida Radiance’, ‘Strawberry Festival’ and ‘Winter Dawn’. Also, RNA-seq libraries from various ‘Camarosa’ tissues^30^ with the study reference PRJEB12420 were downloaded from the European Nucleotide Archive (https://www.ebi.ac.uk/ena). The complete 54 libraries RNA-seq experiment consisted of six independent green receptacle libraries, six white receptacle libraries, six turning receptacle libraries, six red receptacle libraries, three root libraries, three leaf libraries, and six achene libraries each for all corresponding fruit stages. For both libraries, raw RNA-seq reads were assembled to the ‘Camarosa’ reference genome using CLC Genomic Workbench 11 (mismatch cost of 2, insertion cost of 3, deletion cost of 3, length fraction of 0.8, similarity fraction of 0.8, 1 maximum hit per read). Reads that mapped equally well to more than one locus were discarded from the analysis. RNA-seq counts were quantified in Transcripts Per Million (TPM).

### Plant materials for eQTL analysis

Three pedigree-connected and segregating strawberry populations were created from crosses ‘Florida Elyana’ × ‘Mara de Bois’, ‘Florida Radiance’ × ‘Mara des Bois’, and ‘Strawberry Festival’ × ‘Winter Dawn’ (Figure S7). These cultivars and 54 progenies were selected for RNA-seq and SNP genotyping analysis and were used to identify eQTL.

### Fruit transcriptomes analysis of strawberry *MLO* genes for eQTL analysis

Sixty-one fruit transcriptomes were sequenced via Illumina paired-end RNA-seq (Avg. 65million reads, 2×100bp), and consisted of parents and progeny from crosses of ‘Florida Elyana’ × ‘Mara de Bois’, ‘Florida Radiance’ × ‘Mara des Bois’, and ‘Strawberry Festival’ × ‘Winter Dawn’. Reads were trimmed and mapped to the F. ×ananassa octoploid ‘Camarosa’ annotated genome using CLC Genomic Workbench 11 (mismatch cost of 2, insertion cost of 3, deletion cost of 3, length fraction of 0.8, similarity fraction of 0.8, 1 maximum hit per read). Reads that mapped equally well to more than one locus were discarded from the analysis. RNA-seq counts were calculated in Transcripts Per Million (TPM). Three-dimensional principle component analysis (PCA) was performed on all RNA-seq assemblies, including two replicates of ‘Mara des Bois’ fruit harvested three years apart and sequenced independently (Figure S3). Transcript abundances were normalized via the Box-Cox transformation algorithm performed in R^49^ prior to eQTL analysis. The BLAST2GO pipeline was used to annotate the full ‘Camarosa’ predicted gene complement.

### Genome-wide association study (GWAS) of strawberry *MLO* gene expression in fruit

The Affymetrix IStraw35 Axiom® SNP array^50^ was used to genotype 60 individuals, including six parental lines from three independent biparental RNAseq populations. Sequence variants belonging to the Poly High Resolution (PHR) and No Minor Homozygote (NMH) marker classes were included for association mapping. Mono High Resolution (MHR), Off-Target Variant (OTV), Call Rate Below Threshold (CRBT), and Other marker quality classes, were discarded and not used for mapping. Individual marker calls inconsistent with Mendelian inheritance from parental lines were removed. The *F. vesca* physical map was used to orient marker positions as current octoploid maps do not include a majority of the available IStraw35 markers. A genome-wide analysis study (GWAS) was performed using GAPIT v2^51^ performed in R^49^. *MLO* eQTL were evaluated for significance based on the presence of multiple co-locating markers of p-value < 0.05 after false discovery rate correction for multiple comparisons. *MLO* genes demonstrating evidence of an eQTL were then associated using the ‘Holiday’ × ‘Korona’ and family ‘14.95’ octoploid genetic maps to evaluate subgenomic location. Linkage Group in the family ‘14.95’ map was corresponded to the ‘Camarosa’ genome physical chromosomes (Michael Hardigan, UCD, personal communication). Cis vs trans eQTL determinations were made by corroborating known ‘Camarosa’ physical gene position with marker positions in the’14.95’ map, and by BLAST of eQTL markers to the ‘Camarosa’ genome.

## Supporting information

Main Table1

## Acknowledgement

This work was supported by the Florida Strawberry Research and Education Foundation (FSREF), And the “Next-generation Disease Resistance Breeding and Management Solutions for Strawberry” under award no. 2017-51181-26833). We thank all the members of the strawberry genetics and breeding program, and the strawberry molecular genetics and genomics program for their technical assistance.

**Author contributions**

## References Uncategorized References

1. Peries, O.S. Studies on strawberry mildew, caused by Sphaerotheca macularis (Wallr. ex Fries) Jaczewski*. Annals of Applied Biology 50, 211–224 (1962).

2. Xiao, C.L. et al. Comparison of Epidemics of Botrytis Fruit Rot and Powdery Mildew of Strawberry in Large Plastic Tunnel and Field Production Systems. Plant Dis 85, 901–909 (2001).

3. Paulus, A. Fungal Disease of Strawberry. HortScience 25(1990).

4. Peres, N.A., Mertely. J.C. Powdery Mildew of Strawberries. Electronic Data Information Source UF/IFAS Extension (2018).

5. Nelson M.D., G.W.D., Shaw D.V. Relative Resistance of 47 Strawberry Cultivars to Powdery Mildew in California Greenhouse and Field Environments. Plant Disease 80(1996).

6. Devoto, A., Piffanelli, P., Nilsson, I., Wallin, E. Panstruga, R., & Heijne, G.V., Schulze-Lefert, P. Topology, Subcellular, Localization, and Sequence Diversity of the Mlo Family in Plants. The Journal of Biological Chemistry 274, 13 (1999).

7. Kim, M.C. et al. Mlo, a modulator of plant defense and cell death, is a novel calmodulin-binding protein. Isolation and characterization of a rice Mlo homologue. J Biol Chem 277, 19304–14 (2002).

8. Kusch, S. & Panstruga, R. mlo-Based Resistance: An Apparently Universal “Weapon” to Defeat Powdery Mildew Disease. Mol Plant Microbe Interact 30, 179–189 (2017).

9. Pastruga, R. Serpentine plant MLO proteins as entry portals for powdery mildew fungi. Plant Signalling from Genes to Biochemistry 33(2005).

10. Jørgensen, I.H. Discovery, characterization and exploitation of Mlo powdery mildew resistance in barley. Euphytica 63, 141–152 (1992).

11. Acevedo-Garcia, J. et al. The powdery mildew-resistant Arabidopsis mlo2 mlo6 mlo12 triple mutant displays altered infection phenotypes with diverse types of phytopathogens. Sci Rep 7, 9319 (2017).

12. Devoto, A. et al. Molecular phylogeny and evolution of the plant-specific seven-transmembrane MLO family. J Mol Evol 56, 77–88 (2003).

13. Acevedo-Garcia, J., Kusch, S. & Panstruga, R. Magical mystery tour: MLO proteins in plant immunity and beyond. New Phytol 204, 273–81 (2014).

14. Acevedo-Garcia, J. et al. mlo-based powdery mildew resistance in hexaploid bread wheat generated by a non-transgenic TILLING approach. Plant Biotechnol J 15, 367–378 (2017).

15. Zheng, Z. et al. Loss of function in Mlo orthologs reduces susceptibility of pepper and tomato to powdery mildew disease caused by Leveillula taurica. PLoS One 8, e70723 (2013).

16. Nekrasov, V. et al. Rapid generation of a transgene-free powdery mildew resistant tomato by genome deletion. Sci Rep 7, 482 (2017).

17. Njuguna, W., Liston, A., Cronn, R., Ashman, T.-L. & Bassil, N. Insights into phylogeny, sex function and age of Fragaria based on whole chloroplast genome sequencing. Molecular Phylogenetics and Evolution 66, 17–29 (2013).

18. Shulaev, V. et al. The genome of woodland strawberry (Fragaria vesca). Nat Genet 43, 109–16 (2011).

19. Anciro, A. et al. FaRCg1: a quantitative trait locus conferring resistance to Colletotrichum crown rot caused by Colletotrichum gloeosporioides in octoploid strawberry. Theor Appl Genet 131, 2167–2177 (2018).

20. Mangandi, J. et al. Pedigree-Based Analysis in a Multiparental Population of Octoploid Strawberry Reveals QTL Alleles Conferring Resistance to Phytophthora cactorum. G3 (Bethesda, Md.) 7, 1707–1719 (2017).

21. Salinas, N., Verma, S., Peres, N. & Whitaker, V.M. FaRCa1: a major subgenome-specific locus conferring resistance to Colletotrichum acutatum in strawberry. Theor Appl Genet 132, 1109–1120 (2019).

22. Roach, J.A. et al. FaRXf1: a locus conferring resistance to angular leaf spot caused by Xanthomonas fragariae in octoploid strawberry. Theor Appl Genet 129, 1191–201 (2016).

23. Edger, P.P. et al. Origin and evolution of the octoploid strawberry genome. Nat Genet 51, 541–547 (2019).

24. Edger, P.P. et al. Single-molecule sequencing and optical mapping yields an improved genome of woodland strawberry (Fragaria vesca) with chromosome-scale contiguity. Gigascience 7, 1–7 (2018).

25. Miao, L.X. et al. Genomic identification, phylogeny, and expression analysis of MLO genes involved in susceptibility to powdery mildew in Fragaria vesca. Genet Mol Res 15(2016).

26. Dobson, L., Reményi, I. & Tusnády, G.E. CCTOP: a Consensus Constrained TOPology prediction web server. Nucleic acids research 43, W408–W412 (2015).

27. Horton, P. et al. WoLF PSORT: protein localization predictor. Nucleic acids research 35, W585–W587 (2007).

28. Panstruga, R. Discovery of Novel Conserved Peptide Domains by Ortholog Comparison within Plant Multi-Protein Families. Plant Molecular Biology 59, 485–500 (2005).

29. Nguyen, V.N.T., Vo, K.T.X., Park, H., Jeon, J.-S. & Jung, K.-H. A Systematic View of the MLO Family in Rice Suggests Their Novel Roles in Morphological Development, Diurnal Responses, the Light-Signaling Pathway, and Various Stress Responses. Frontiers in Plant Science 7(2016).

30. Sánchez-Sevilla, J.F. et al. Gene expression atlas of fruit ripening and transcriptome assembly from RNA-seq data in octoploid strawberry (FragarialJ×lJananassa). Scientific Reports 7, 13737 (2017).

31. Jiao, Y. & Paterson, A.H. Polyploidy-associated genome modifications during land plant evolution. Philosophical transactions of the Royal Society of London. Series B, Biological sciences 369, 20130355 (2014).

32. Tao, Y., Zhao, X., Mace, E., Henry, R. & Jordan, D. Exploring and Exploiting Pan-genomics for Crop Improvement. Mol Plant 12, 156–169 (2019).

33. Mutti, J.S., Bhullar, R.K. & Gill, K.S. Evolution of Gene Expression Balance Among Homeologs of Natural Polyploids. G3 (Bethesda) 7, 1225–1237 (2017).

34. Chen, Z. et al. Expression analysis of the AtMLO Gene Family Encoding Plant-Specific Seven-Transmembrane Domain Proteins. Plant Molecular Biology 60, 583–597 (2006).

35. Pessina, S. et al. Characterization of the MLO gene family in Rosaceae and gene expression analysis in Malus domestica. BMC Genomics 15, 618 (2014).

36. Jiwan, D., Roalson, E.H., Main, D. & Dhingra, A. Antisense expression of peach mildew resistance locus O (PpMlo1) gene confers cross-species resistance to powdery mildew in Fragaria x ananassa. Transgenic Res 22, 1119–31 (2013).

37. Consonni, C. et al. Conserved requirement for a plant host cell protein in powdery mildew pathogenesis. Nat Genet 38, 716–20 (2006).

38. Leister, D. Tandem and segmental gene duplication and recombination in the evolution of plant disease resistance gene. Trends Genet 20, 116–22 (2004).

39. Panchy, N., Lehti-Shiu, M. & Shiu, S.H. Evolution of Gene Duplication in Plants. Plant Physiol 171, 2294–316 (2016).

40. Yang, T., Peng, H. & Bauchan, G.R. Functional analysis of tomato calmodulin gene family during fruit development and ripening. Horticulture research 1, 14057–14057 (2014).

41. Voorrips, R.E. MapChart: Software for the Graphical Presentation of Linkage Maps and QTLs. Journal of Heredity 93, 77–78 (2002).

42. Guo, A.-Y. et al. GSDS 2.0: an upgraded gene feature visualization server. Bioinformatics 31, 1296–1297 (2014).

43. Kearse, M. et al. Geneious Basic: an integrated and extendable desktop software platform for the organization and analysis of sequence data. Bioinformatics (Oxford, England) 28, 1647–1649 (2012).

44. Sarkar, D. Lattice: Multivariate Data Visualization with R. Springer (2008).

45. Brors, B., Gu, L., Schlesner, M., Eils, R. & Gu, Z. circlize implements and enhances circular visualization in R. Bioinformatics 30, 2811–2812 (2014).

46. Marchler-Bauer, A. et al. CDD: a Conserved Domain Database for the functional annotation of proteins. Nucleic acids research 39, D225–D229 (2011).

47. Liu, W. et al. IBS: an illustrator for the presentation and visualization of biological sequences. Bioinformatics 31, 3359–61 (2015).

48. Mitchell, C. MultAlin–multiple sequence alignment. Bioinformatics 9, 614–614 (1993).

49. Team, R.D.C. R: A Language and Environment for Statistical Computing. R Foundation for Statistical Computing, Vienna, Austria (2014).

50. Verma, S. et al. Development and evaluation of the Axiom® IStraw35 384HT array for the allo-octoploid cultivated strawberry Fragaria ×ananassa. 1156 edn 75–82 (International Society for Horticultural Science (ISHS), Leuven, Belgium, 2017).

51. Tang, Y. et al. GAPIT Version 2: An Enhanced Integrated Tool for Genomic Association and Prediction. The Plant Genome 9(2016).

