## Supplemental Figure Legend for "Genome-Wide Identification and Characterization of *MLO* Gene Family in Octoploid Strawberry (*Fragaria ×ananassa*)"

**Genome-Wide Identification and Characterization of *MLO* Genes Associated with Strawberry (*Fragaria ×ananassa*) Powdery Mildew and Fruit Ripening**

Ronald R. Tapia^1^, Christopher R. Barbey^2^ Saket Chandra^2^, Kevin M. Folta^2^, Vance M. Whitaker^1^ and Seonghee Lee^1†^

^1^Department of Horticultural Sciences, University of Florida, IFAS Gulf Coast Research and Education Center, Wimauma, FL 33598, USA.

^2^Department of Horticultural Sciences, University of Florida, 1301 Fifield Hall, PO Box 110690, Gainesville, FL 32611, USA.

Supplementary Figures S1-S4

Supplementary Tables S1-S4

**Supplementary Figure Legends**

**Figure S1.** Exon-intron structures of strawberry *MLO* genes predicted from *F. vesca* (A) and *F. ×ananassa* (B) genomes. Genetic structures of 20 *FveMLO* and 68 *FaMLO* genes were constructed using Gene Structure Display Server 2.0 (GSDS).

**Figure S2.** Pairwise comparison of putative *MLO* coding DNA sequences (CDS) of *F. vesca* and *F. ×ananassa*. Percent sequence similarities among predicted 20 *FveMLO* and 68 *FaMLO* genes were calculated using multiple sequence alignment and tree building programs via Geneious software. Gene sequences similarities were visualized with heatmap using R-package “lattice”.

**Figure S3.** Multiple protein sequences alignment of strawberry candidate *MLO* proteins and known *AtMLO* proteins. Amino acid sequence of *FveMLO10*, *FveMLO17* and *FveMLO20* from *F. vesca*; *FaMLO10-1*, *FaMLO10-4*, *FaMLO17-1*, *FaMLO17-2*, *FaMLO17-3*, *FaMLO17-4*, *FaMLO20-1*, *FaMLO20-2*, *FaMLO20-3* and *FaMLO20-4* from *F. ×ananassa*; *AtMLO2, AtMLO6* and *AtMLO12* from *A. thaliana* were aligned with MultAlin using default settings. The positions of transmembrane (TM1-TM7) domains^8^, predicted CAMBD^18^ and the other two C-terminal domains (I and II)^7^ were highlighted and indicated with lines.

**Figure S4**. *FaMLO* transcript level distribution for marker genotypes of the most significant SNPs identified in eQTL analysis.

**Supplementary Tables**

**Table S1.** Characteristics of strawberry *MLO* genes. (A) *FveMLO* genes predicted using *F. vesca* (v4.0.a1) reference genome sequence and (B) *FaMLO* genes predicted using *F. ×ananassa* (v1.0.a1) reference genome sequence (https://www.rosaceae.org/)

**Table S3.** Table S3. Topology and Subcellular localization of putative *FveMLO* (A) and *FaMLO* (B) proteins. Transmembrane (TM) and *MLO* domains were predicted using online software CCTOP and CDD:NCBI’s conserved domain database, respectively. Subcellular localization of *MLO* genes was predicted using WoLF PSORT.

**Table S4.** Expression profile of putative strawberry *MLO* genes in three pedigree-connected and segregating strawberry populations including parental genotypes, ‘Florida Elyana’, ‘Mara de Bois’, ‘Florida Radiance’, ‘Strawberry Festival’ and ‘Winter Dawn’. Total RNA was isolated from mature fruit tissues followed by cDNA library construction. Then, samples were sent for sequencing using Illumina Hi-Seq 4000. RNA-seq data was assembled and annotated using the recently published *F. ×ananassa* cv ‘Camarosa’ reference genome. The raw gene expression level was quantified using Transcripts Per Million (TPM) as unit.
