## Supplemental Figures for "Genome-Wide Identification and Characterization of *MLO* Gene Family in Octoploid Strawberry (*Fragaria ×ananassa*)"

### Slide 1
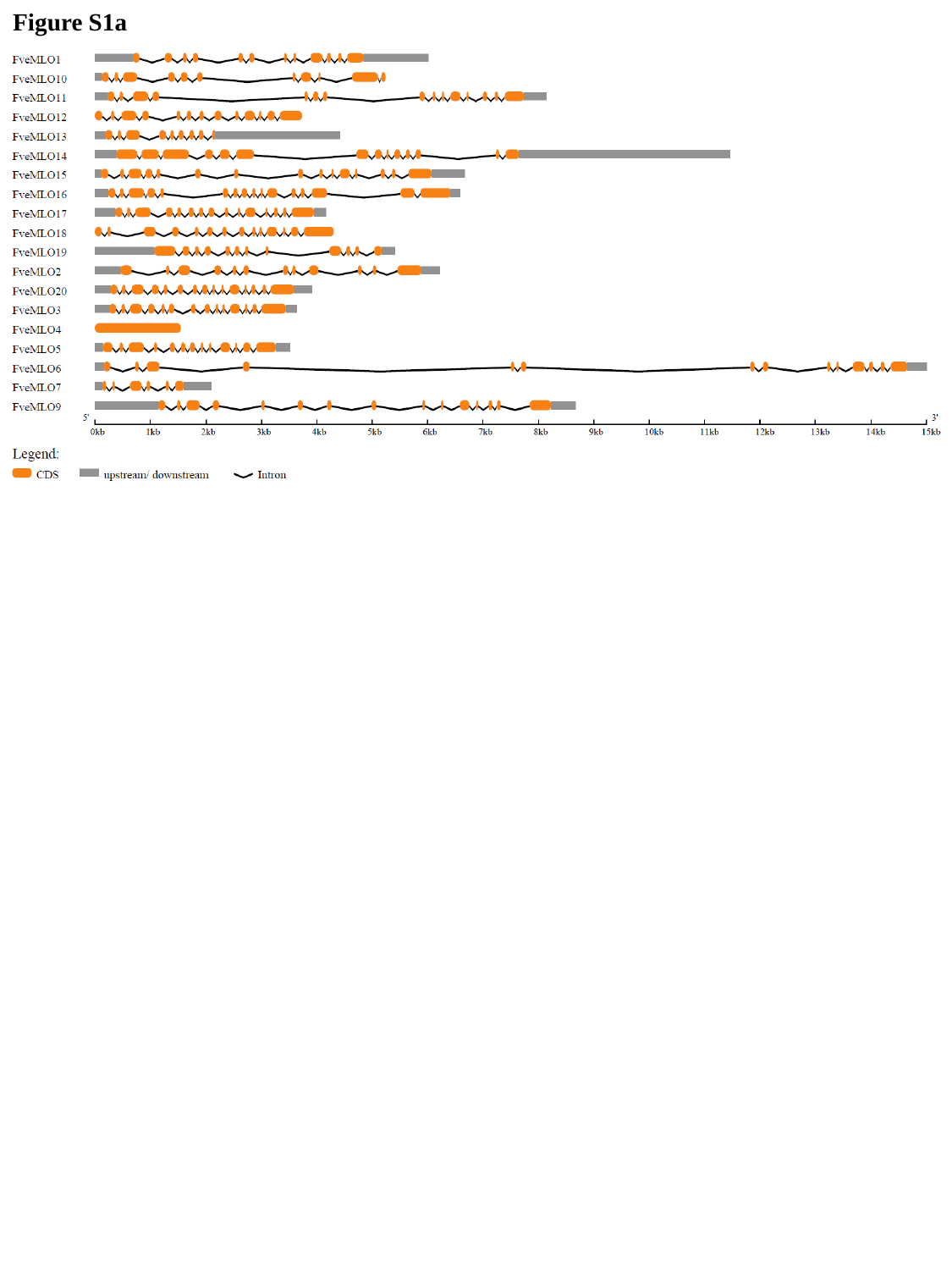

Figure S1a

### Slide 2
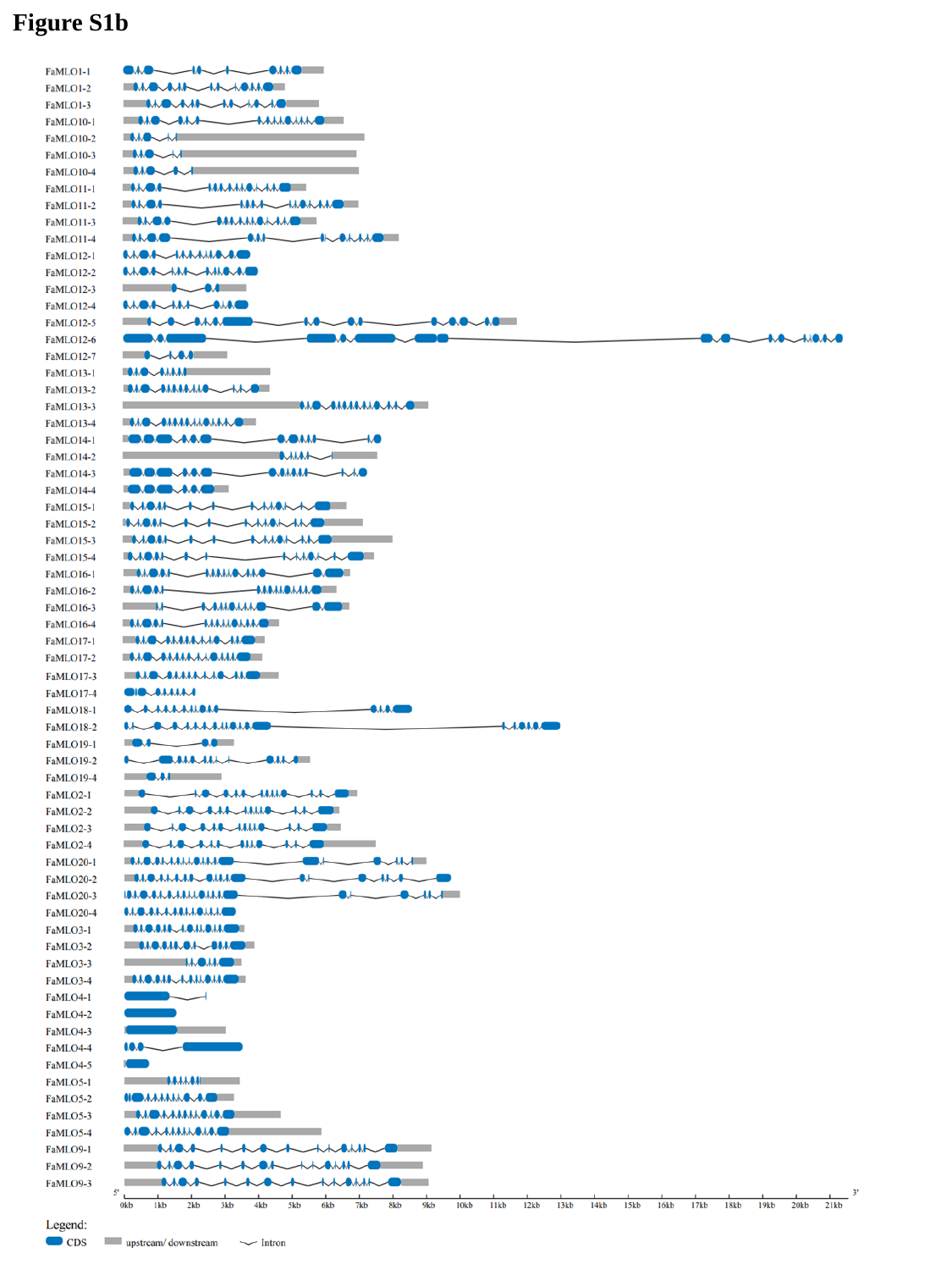

Figure S1b

### Slide 3
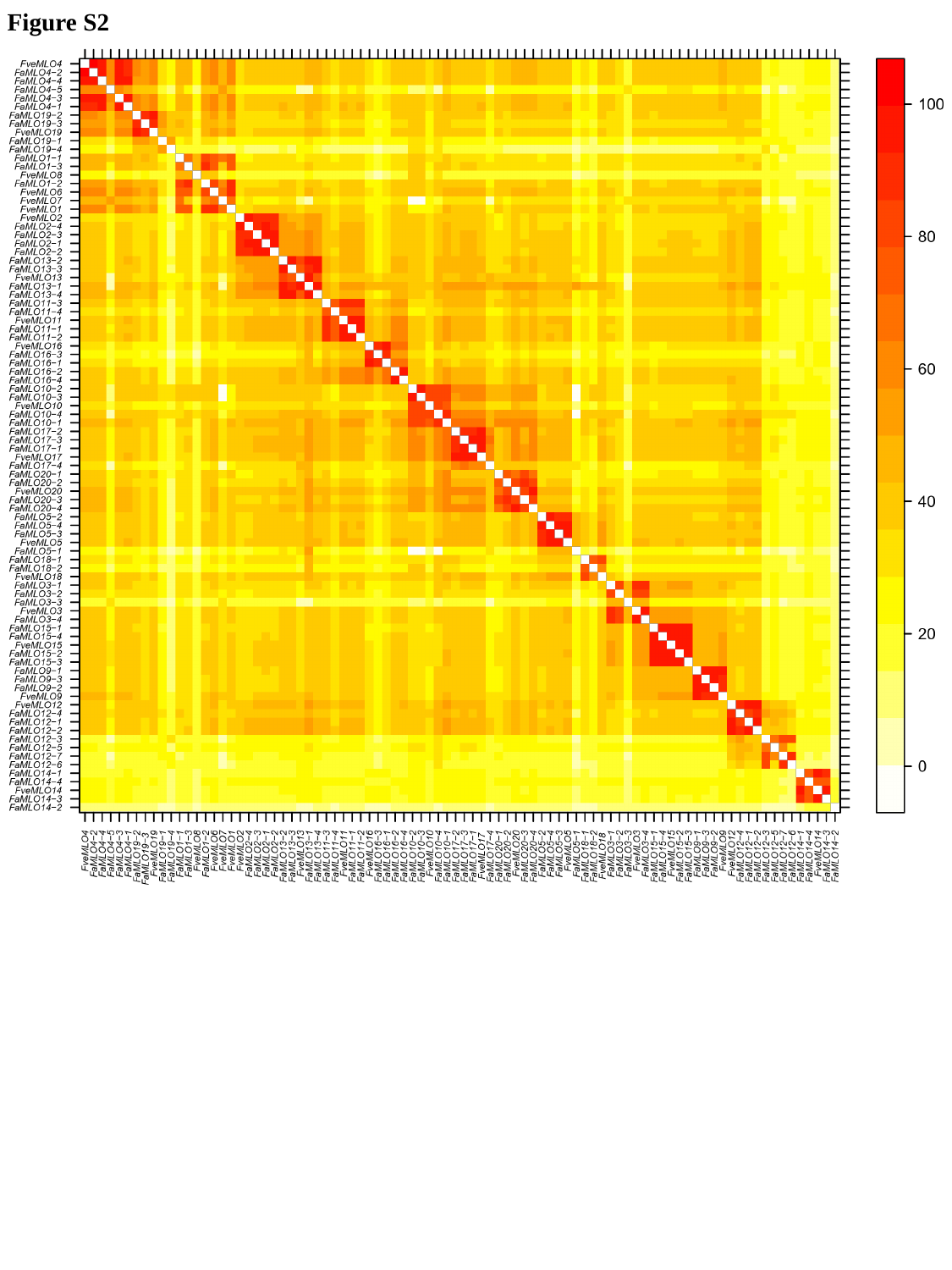

Figure S2

### Slide 4
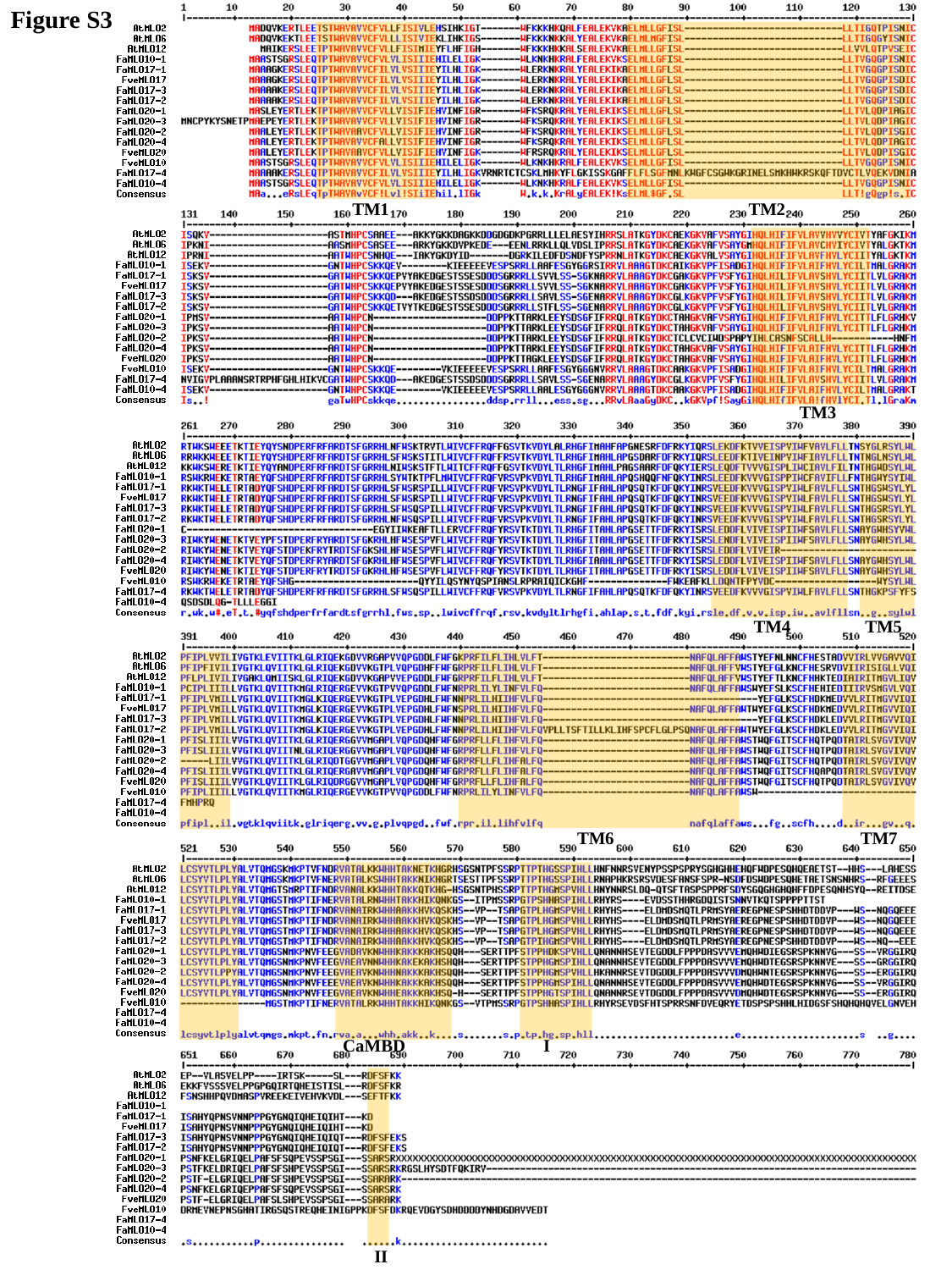

Figure S3
TM2
TM1
TM3
TM4
TM5
TM6
TM7
I
CaMBD
II

### Slide 5
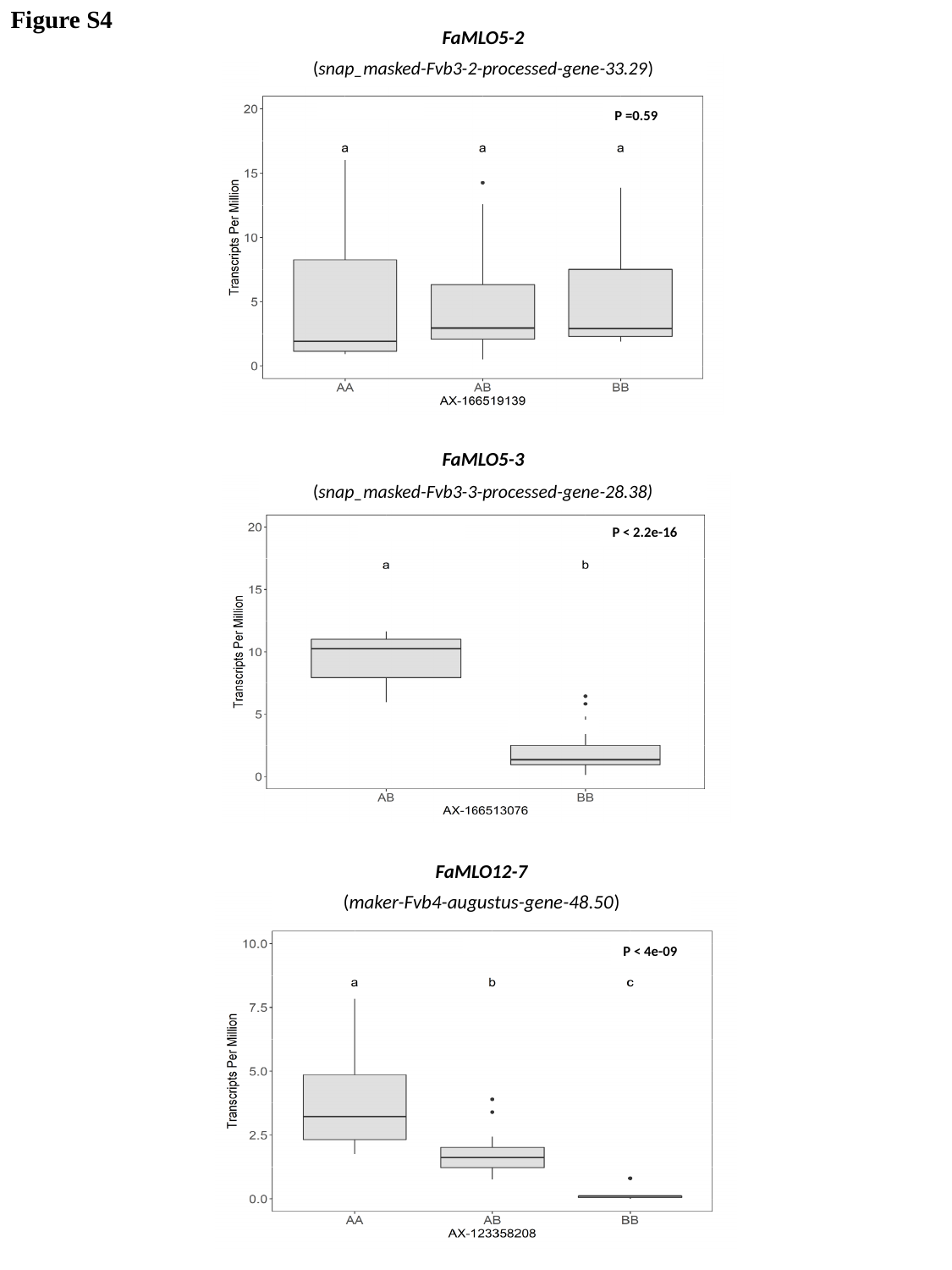

Figure S4
FaMLO5-2
(snap_masked-Fvb3-2-processed-gene-33.29)
P =0.59
FaMLO5-3
(snap_masked-Fvb3-3-processed-gene-28.38)
P < 2.2e-16
FaMLO12-7
(maker-Fvb4-augustus-gene-48.50)
FaMLO12-7
P < 4e-09
